# Trophic specialization enhances growth performance in larvae of southern bluefin, albacore, and skipjack tunas from the eastern Indian Ocean

**DOI:** 10.1101/2025.08.28.672816

**Authors:** Raúl Laiz-Carrión, Ricardo Borrego-Santos, José María Quintanilla, Claudio Quezada-Romegialli, Estrella Malca, Rasmus Swalethorp, Francisco Abascal, María Grazia Pennino, Manuel Vargas-Yáñez, Miguel Ángel Godoy-Bermúdez, David Die, Michael R. Landry

## Abstract

We examined trophic ecology and its influence on larval growth variability for three scombrids, southern bluefin (*Thunnus maccoyii*, SBT), albacore (*T. alalunga*, ALB), and skipjack tunas (*Katsuwonus pelamis*, SKJ), that share a common spawning ground in the eastern Indian Ocean. We combined otolith-based ageing with bulk nitrogen and carbon stable isotope analysis (SIA) of individual larvae. Significant interspecific differences in δ¹⁵N and δ¹³C indicate adaptive resource partitioning that allows these tunas to coexist during early ontogeny. Trophic position and isotopic niche were estimated with both frequentist and Bayesian approaches, enabling the evaluation of ontogenetic isotope shifts, niche overlap, and resource use in relation to growth. ALB grew fastest had the highest trophic position, and the broadest isotopic niche. Optimally growing tuna larvae occupied the narrowest trophic niche and had lower trophic positions for all three species, supporting the hypothesis that strong trophic specialization supports better growth performance, and that feeding on more efficient shorter food chains (e.g., microbial loop via appendicularians) can enhance larval fitness. Using lower C:N ratio as a proxy of larval condition, found in optimal growing groups, supports the broader hypothesis that growth potential is closely tied to energy allocation strategies during early ontogeny. A detailed understanding of how larval trophodynamics, niche breadth, and resource partitioning interact with growth and survival during these vulnerable stages is essential for ecosystem-based management, particularly in systems where growth rate modulates predation risk and competitive fitness.

## 1. Introduction

The tunas (tribe Thunnini) include the most economically important group of species, known as the principal market tunas, notably southern bluefin tuna (*Thunnus maccoyii*, SBT), albacore tuna (*Thunnus alalunga*, ALB), and skipjack tuna (*Katsuwonus pelamis*, SKJ), along with other highly valuable tunas species (see Muhling et al., 2017). These tuna species are top predators that play a significant role in open-ocean ecosystems, due to their influence on the structure and dynamics of marine food webs (Estes et al., 2016; van Denderen et al., 2018). Hence, their decline can initiate trophic cascades (Heithaus et al., 2008) and endanger the resilience and stability of marine resources (Kerr et al., 2017).

SBT is one of the largest, longest-lived and longest-migrating tunas and represents a commercially valuable species distributed across temperate waters of the Southern Hemisphere (Joan-Jordá et al., 2013). Its range spans the eastern Atlantic, Indian, and southwestern Pacific Oceans, typically between 30°S and 50°S (Caton, 1991). Unlike tropical tunas that spawn year-round across broader geographic ranges, SBT undertake long-distance migrations to a single, spatially restricted spawning ground overlying the 5000-m deep Argo Abyssal Plain (hereafter, Argo Basin) in the tropical northeastern Indian Ocean (IO) between Indonesia and northwestern Australia, where spawning is limited to a brief seasonal window (Farley and Davis, 1998; Patterson et al., 2008; Farley et al., 2015; Hobday et al. 2015). Genetic analyses confirm the existence of a single, panmictic population (Grewe et al., 1997) considered to constitute a heavily overfished single stock for management purposes (Annon, 2021). However, overfishing is not currently occurring due to measures taken in a rebuilding plan (ISSF 2025) by the Commission for the Conservation of Southern Bluefin Tuna (CCSBT). Compared to other bluefin tuna species, only limited information on SBT spawning habitat, larval ecology, and behavior is available from surveys conducted several decades ago (Nishikawa 1985, Nishikawa and Rimmer, 1987, Davis et al., 1990a, 1990b, Jenkins and Davis, 1990).

ALB is a widely distributed and highly migratory species occurring in all tropical, subtropical, and temperate pelagic ecosystems (Essington, 2003) with seasonal movements influenced by oceanographic conditions (Nikolic et al., 2017). ALB spawn seasonally in IO subtropical waters during the southern summer months (October–March) when water temperatures exceed 24°C (Nishikawa, 1985). In the eastern IO, potential spawning areas are mostly between 10 and 30° latitude, including regions north of Australia, around Indonesia, and the eastern boundary currents where warm oligotrophic waters prevail (Nishikawa, 1985; Reglero et al., 2014). The Indian Ocean albacore stock is not overfished and is not subject to overfishing. However, there is considerable uncertainty associated with the latest stock assessment (ISSF 2025).

SKJ is also a highly migratory species with a widespread presence across the tropical and subtropical waters of three major oceans (Atlantic, Indian, and Pacific) (Artetxe-Arrate et al., 2021). SKJ spawn year-round in equatorial and tropical waters, and seasonally in subtropical waters when surface water temperature exceeds 24°C (Nishikawa, 1985). In Indonesia waters, spawning is is driven by changes in oceanic conditions with peaks coinciding with seasonal upwelling driven by the Southeast Monsoon occurring between June through September. Climate oscillations such as the Indian Ocean Dipole (IOD) (Marsac and Le Blanc, 1998) and El Niño-Southern Oscillation (ENSO) influence spawning success, larval survival, and recruitment (Marsac and Le Blanc, 1999). SKJ dominated tuna catches in the Indian Ocean, where tunas and tuna-like species are the main contributors to the total marine catch (FAO, 2024). Recent assessments report no evidence of overfishing, and no conservation measures have yet been adopted specifically for SKJ tuna (ISSF, 2025).

Historical larval surveys from the mid-1950s to early 1980s delineated the SBT primary spawning area between ∼ 7°S–20°S and 110°E–125°E (Nishikawa, 1985). In this shared spawning ground for SBT, ALB, and SKJ, the upper ocean is characterized by warm oligotrophic conditions, with surface temperatures exceeding 26°C, reduced salinity (∼34), strong stratification, low eddy kinetic energy, and low chlorophyll-a concentrations (Nieblas et al., 2014). This region experiences wind-driven vertical mixing during the mid-summer cyclone season, which coincides with the peak of SBT spawning (Davis et al., 2003; Willis and Hobday, 2007; Nieblas et al., 2014; Jiang et al., 2022). These environmental features influence nutrient availability and food-web dynamics. The “loophole hypothesis” (Bakun and Broad, 2003) suggests that while oligotrophic conditions may limit larval food availability, they concurrently reduce predation pressure, potentially enhancing larval survival. Understanding how food resources are partitioned among species to limit competition is crucial for explaining differences in growth potential and survival during these vulnerable larval stages.

Stable isotope analysis (SIA) of nitrogen (δ¹⁵N) and carbon (δ¹³C) provides insights into the sources of energy and nutrients assimilated by consumers, as well as their trophic positions (TPs) within food webs (Peterson and Fry, 1987; Zanden and Rasmussen, 2001; Post, 2002). Traditional trophic studies on tuna larvae have relied on stomach content analysis, providing hours-long snapshots of prey consumption (e.g., Young and Davis 1990; Catalán et al., 2007; Llopiz et al., 2015; Uriarte et al., 2019; Shiroza et al., 2022: Swalethorp et al., this issue). However SIA provides complementary insights reflecting several days of the trophic ecology of fish larvae (Laiz-Carrion et al., 2011, 2022; Quintanilla et al., 2015, 2020), including tuna larvae (Laiz Carrion et al., 2013, 2015, 2019; Malca et al., 2023; García et al., 2017; Quintanilla et al., 2023, 2024). This approach provides time-integrated information, reflecting assimilated rather than ingested food, making it a suitable approach for testing hypotheses regarding developmental shifts in food sources associated with larval growth variability (García et al., 2017; Malca et al., 2022, 2023; Quintanilla et al., 2015, 2020, 2023, 2024). Specifically, δ¹⁵N values have been widely used to estimate the TPs of marine organisms due to the stepwise enrichment of nitrogen isotopes with each trophic transfer (Post, 2002; Parnell et al., 2013; Quezada-Romegialli et al., 2018) with size-fractionated zooplankton serving as baseline indicators (Laiz-Carrión et al., 2015). Meanwhile, δ¹³C values provide insights into the origin of primary production supporting a consumer’s diet, often indicating habitat-specific feeding (France, 1995; Phillips and Gregg, 2001; Burbank et al., 2024). Furthermore, SIA data has been used to quantify isotopic niche width and overlap, providing insights into resource partitioning and the degree of dietary and habitat overlay among species or life stages (Layman et al., 2007; Newsome et al., 2007; Jackson et al., 2011; Parnell et al., 2010; Swanson et al., 2015). These isotopic niches, visualized through bivariate δ¹³C and δ¹⁵N space, can reveal temporal and/or spatial variability in resource use, and help identify the extent of trophic interactions among co-occurring taxa (Layman et al., 2011). In particular, for scombrid larvae, TP estimations and isotopic niche analysis have revealed patterns of resource use, predator-prey dynamics (Polunin et al., 2002), interspecific trophic differentiation, and trophic influences on growth, enhancing our understanding of larval trophic ecology, ecological niches, and food-web dynamics. (Laiz-Carrión et al., 2015).

Larval survival is strongly linked to growth potential (Anderson, 1988). According to the growth dependent mortality hypothesis, feeding modulates growth rates and therefore size of individuals, and eventually predation pressure (e.g., Nielsen and Munk, 2004). The functional basis of this hypothesis was expanded by Hare and Cowen (1997) by proposing three basic concepts that interrelate during larval development: bigger is better (Miller et al., 1988; Leggett and DeBlois, 1994), growth rate (Ware, 1975; Shepherd and Cushing 1980) and the stage-duration hypotheses (Chambers and Leggett, 1987; Houde, 1987). The two principal environmental factors that influence growth variability are temperature and food availability (Buckley, 1984; Heath, 1992). Combining SIA with larval size and estimations of larval age from otolith microstructure analysis allows the evaluation of different growth strategies throughout the ontogenetic developmen and in relation to trophic dynamics.

This study provides a comprehensive understanding of larval trophodynamics on growth variability, possible implications for recruitment, and evaluates nursery habitat quality for three larval tunas (SBT, ALB, and SKJ) that share a common IO spawning ground. First, we assesses the influence of feeding on larval growth variability by comparing daily growth and isotopic signatures of δ¹⁵N and δ¹³C for individual larvae. Next, we evaluate larval TP and trophic niches for each species, assess SIA ontogenetic shifts, trophic niche overlap, and evaluate resource partitioning. Our comparative study provides valuable insights into how these three co-occurring marine top-predators compete and/or partition space along with prey resources during their most vulnerable developmental period, the larval period.

## 2. Materials and methods

### 2.1. Environmental measurements

Hydrographic profiles of the ocean environment were collected at each sampling station using vertical casts with a Seabird SBE 25plus CTD system, mounted on a rosette equipped with 12 Niskin bottles and sensors for measuring temperature (°C), salinity (psu), fluorescence (volts), and dissolved oxygen concentration (mg l^−1^). Environmental characterization focused on the upper 25 m of the water column and included measurements of temperature, salinity, fluorescence, oxygen as well as mixed layer depth (m, hereafter MLD), defined as the depth at which σ_T_ density first exceeded the 0-5 m average by 0.01 kg m^-3^. We include also micro- and meso- zooplankton biomasses (mg/m³) and their N and C stable isotope values. A detailed description of the CTD deployment and sampling protocols is provided in Romero-Fernández et al. (this issue).

### 2.2. Tuna larvae and zooplankton sampling

As part of the 2nd International Indian Ocean Expedition (IIOE-2), tuna larvae and environmental data were collected during BLOOFINZ-IO cruise RR2201 aboard R/V *Roger Relvelle* from January to March 2022, coinciding with the peak SBT spawning period (Fig. 1). Icthyoplankton samples were collected using methodologies previously described for larval studies of Atlantic Bluefin Tuna (ABT) in the Mediterranean Sea and Gulf of Mexico (Laiz-Carrión et al., 2013, 2015, 2019; Malca et al., 2022, 2023; Quintanilla et al., 2024). Larval specimens were collected with double oblique tows from a depth of 25 m to the surface with a 90-cm Bongo net (0.5-mm mesh) and a 1-m² surface net (1-mm mesh), both equipped with General Oceanics flowmeters (details in Landry et al., this issue). SBT, ALB, and SKJ larvae were identified at-sea utilizing morphological, meristic, and pigmentation characteristics (Nishikawa, 1985; Nishikawa and Rimmer 1987). All tuna larvae were subsequently preserved at -80°C for for subsequent morphometrics, genetics, SIA, and ageing in the IEO-CSIC laboratories. Shipboard identifications were confirmed using a multiplex PCR to distinguish SBT from yellowfin (*T. albacares*), albacore (*T. alalunga*), bigeye (*T. obesus*), and skipjack (*K. pelamis*) tunas (details in Malca et al., this issue).

**Figure 1.**
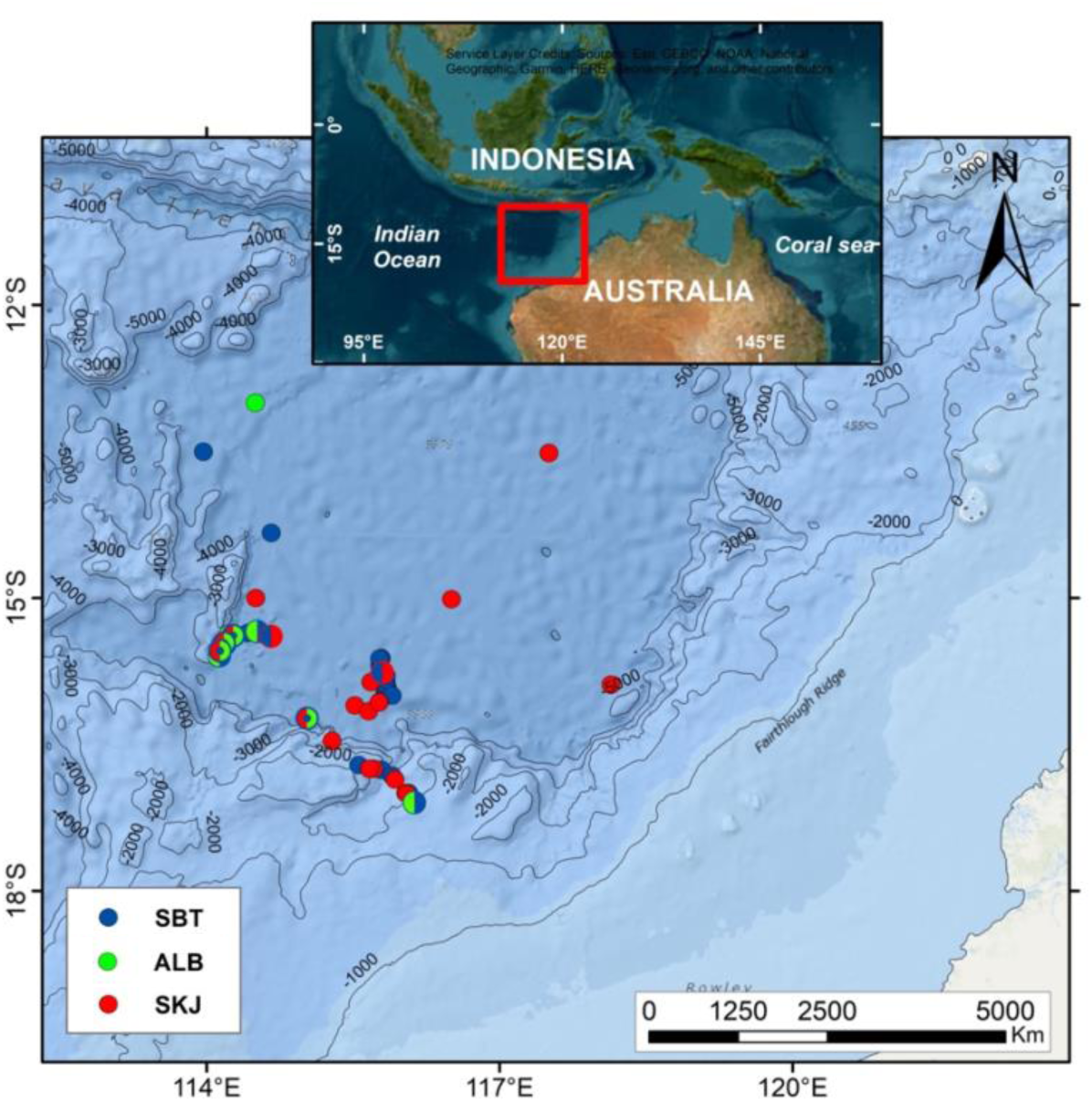
General study area for BLOOFINZ-IO in the Eastern Indian Ocean (IO) during the 2022 spawning season, showing station locations of scombrid larvae analyzed for stable isotope analysis. Different colors represent the different species analyzed: blue for *Thunnus maccoyii* (SBT), green for *T. alalunga* (ALB) and red for *Katsuwonus pelamis* (SKJ).

Coupled to the Bongo-90 net system, a Bongo-25 (25 cm diameter) plankton net was fitted with 200 and 55 μm mesh nets to simultaneously target meso- and micro-zooplankton, respectively. A mechanical flowmeter (2030, General Oceanics) was placed in the center of all plankton nets to calculate the volume of water filtered (m^3^) by each net. Plankton samples from the 200-μm net (hereafter meso-zooplankton) were divided into 2 aliquots using a Folsom plankton splitter, with half preserved in 95% ethanol and the other half frozen at −80°C. Samples from the 55-μm mesh net (hereafter micro-zooplankton) were first poured through a 200-μm mesh sieve to remove larger zooplankton, then filtered onto 55-μm mesh and frozen at −80°C. Each meso- and micro-zooplankton sample was freeze-dried for 48 h at −20°C, and weighed to the nearest 1 μg. Dry weight (DW) biomass values were standardized to mg m^−3^ using the volume filtered by the plankton nets.

### 2.3. Tuna larval processing

In the laboratory, the larvae were measured for standard length (SL, mm) with Image J 1.44a software (USA National Institute of Health) and dry weights (DW, mg) were recorded to the nearest 1 μg after freeze-drying for 24 h. A total of 463 larvae (251 SBT, 156 ALB and 56 SKJ) within a 4–9 mm size range were selected for DW vs SL relationship analysis.

Larvae were classified as preflexion and postflexion (Kendall et al., 1984) based on the degree of notochord flexion observed on a digitized and calibrated image of each larva (see Borrego-Santos et al., this issue).

Next, otoliths were removed andage estimations were conducted following the protocols described by Borrego-Santos et al., *this issue*. Individual tuna larvae were grouped by their positive and negative residuals of SL-at-age and DW-at AGE, considered simultaneously. The fastest growing or optimally growing (OPT) had positive residuals for SL-at-age and DW-at-age while the slowest growing or deficient (DEF) tunas had negative residuals for both metrics (Quintanilla et al., 2015; details in Borrego-Santos et al., this issue).

### 2.4. Stable isotope analysis (SIA)

Individual larvae that were prepared for ageing were also processed for SIA. 307 larvae (163 SBT, 90 ALB, and 53 SKJ) that were within the 4–9 mm common size range were selected for age and SIA analyses. Although most larvae (particularly SBT) were collected at night, when they were unlikely to be feeding, their stomachs were removed before the remaining body tissue was placed in a 0.03-ml tin capsule.

Natural abundances of N (δ^15^N) and C (δ^13^C) were measured using an isotope-ratio spectrometer (Thermo-Finnigan Delta-plus) coupled to an elemental analyzer (FlashEA1112 Thermo-Finnigan) at the Instrumental Unit of Analysis of the University of A Coruña. Ratios of ^15^N/^14^N and ^12^C/^13^C were expressed in conventional delta notation (δ), relative to the international standard, Atmospheric Air (N_2_) and Pee-Dee Belemnite (PDB) respectively, using acetanilide as standard. The analysis precisions for δ^15^N and δ^13^C were 0.11‰ and 0.14‰, respectively, based on the standard deviation of internal references (repeatability of duplicates). Prior chemical extraction for lipid correction was not possible to carry out due to the low amount of sample available. Nevertheless, a posterior lipid content correction of the δ^13^C values was conducted following the model proposed by Logan et al. (2008) for averaged ABT muscle tissue, obtaining a mean ± SD values of 8.59 ± 1.08‰, 8.78 ± 1.14‰ and 8.72 ± 1.32‰ lipid correction for SBT, ALB and SKJ larvae respectively. This procedure has been previously described for scombrid larvae by Laiz-Carrión et al. (2015). C:N ratios were determined as elemental ratios.

### 2.5. Isotopic niche widths

Only postflexion larvae were selected for isotopic niche widths and TP analyses. Isotopic niche widths were estimated both by standard Bayesian ellipse areas and associated credible interval adjusted for small sample size (SEAc) (Jackson et al., 2011, 2012) and by Kernel utilization density (KUD; rKIN package; Eckrich et al., 2020). Isotopic niche widths and overlap analyses were conducted using the R package SIBER (Stable Isotope Bayesian Ellipses in R) v.3.3.0 (Jackson et al., 2011, R Core Team, 2024). The standard ellipses for bivariate data were calculated from the variance and 40% covariance of the data following Laiz-Carrión et al. (2019). The KUD (40% contour) method is less sensitive to extreme values and low number of samples; in order to incorporate sampling uncertainty, we applied a non-parametric bootstrap procedure with n = 1000 iterations for each species (SBT, ALB, SKJ) and physiological growth condition (OPT, DEF). In each iteration, a random sample with replacement was drawn from the original data, preserving its sample size. The 95% kernel density area was then estimated using the estKIN function. Iterations were retained only when the isotopic niche estimation was successful (i.e., at least 3 unique data points and successful convergence of the algorithm). The resulting bootstrap distributions of SEA values were used to compute summary statistics (mean, median, 95% confidence interval) and to visualize variability within each group using violin plots with overlaid boxplots, means and medians. For both SEA_C_ and KUD, overlap was estimated as the ratio of overlapped area to total area for each species pair comparison

### 2.6. Trophic position estimation

TPs were estimated only for postflexion larvae to avoid maternal influence on the δ^15^N values (Uriarte et al., 2016; Laiz-Carrión et al., 2019), according to Equation (1) applying frequentist (Post, 2002) and Bayesian statistical (Quezada-Romegialli et al., 2018; 2024) inferences.

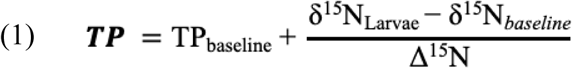

In Equation 1, δ^15^N_larva_ represents the isotopic value of individual larvae and δ^15^N_baseline_ is the microzooplankton isotopic values at the same station where the larva was collected. TP_baseline_ is the baseline consumer trophic position represented by the microzooplankton (0.05 – 0.2 mm), which consists of primary producers and primary consumers with a designated value of 2 (Coll et al., 2006; Bode et al., 2007; Laiz-Carrión et al., 2015; Quezada-Romegialli et al., 2018; Laiz-Carrión et al., 2019; Malca et al., 2023; Quintanilla et al., 2024). The Δ^15^N is an experimental nitrogen isotopic discrimination factor with a mean value of 1.46 ‰, (± 0.2 as standard deviation, SD) as proposed by Varela et al. (2012) for ABT juveniles.

For the Bayesian framework, trophic position (TP) was estimated using the tRophicPosition package, which implements Markov Chain Monte Carlo (MCMC) simulations to model uncertainty in isotopic data (Quezada-Romegialli et al., 2018). This approach incorporates variability in consumer and baseline isotopic values, as well as in the trophic discrimination factor. Specifically, we applied the *oneBaseline* model, which assumes a single isotopic baseline, and adapted it for individual-level TP estimation. While tRophicPosition v0.8.0 only provides population-level estimates, we developed a custom implementation of the *oneBaseline* model using the *greta* framework (Golding, 2019), enabling individual-level inference as recently demonstrated by Quezada-Romegialli et al. (2024). Briefly, each baseline was modeled using a normal distribution defined by the mean of δ^15^N_baseline_ ± 0.5 (SD), with SD modeled with a Cauchy distribution (scale = 3), truncated to positive values. Baseline trophic level was fixed as 2 and the trophic discrimination factor (TDF) was modeled as a normal distribution with a mean of 1.46 ‰ and a standard deviation of 0.16, following Varela et al. (2012). The analysis was run with 1,000 samples as warmup and 10,000 posterior samples, a thinning of 10 and 16 parallel Markov chains. Figures were done using the tidybayes v3.06 (Kay, 2023), tidyverse (Wickham et al., 2019) and patchwork v1.3.0 (Pedersen, 2024) packages in R v4.3.0 (R Core Team, 2024) within RStudio v2025.5.1.513 (Posit team, 2025).

### 2.7. Statistical analyses

To assess environmental variability and potential trophic influences on larval growth, the measured variables compared using a non-parametric Kruskal-Wallis test across species (SBT, ALB and SKJ) and the Mann-Whitney U test between OPT and DEF groups within each species, which did not fit parametric assumptions. Least squares linear regression was applied to SL and DW as functions of AGE to characterize the daily growth patterns of each species. Analyses of covariance (ANCOVA) were run for SL with age as a covariate and for growth rate (mm d^-1^) differences and trophic variables among the three species. In addition, ANCOVA was conducted between OPT and DEF groups using AGE as the covariate within species. Variables were log-transformed (LOG) prior to statistical analysis when necessary to obtain linearity and variance homogeneity (Sokal and Rohlf, 1979). To evaluate the relationship between individual trophic position (TP) and ontogeny, we fitted two polynomial models incorporating age (in days) as a second-degree polynomial predictor. First, to assess species-specific differences in TP trajectories with age independently of nutritional condition, we fitted a global model specified as TP ∼ SPP × poly(AGE, 2), where SPP is a categorical variable representing species identity (SBT, ALB, SKJ), and poly (AGE, 2) captures potential non-linear relationships between age and individual TP. This approach allowed us to test whether both the intercepts and the linear and quadratic components of the individual TP-age relationship differed among species. Second, to explore the effects of nutritional condition (OPT vs DEF) within each species, we fitted species-specific models defined as TP ∼ OPT-DEF × poly(AGE, 2), where OPT-DEF represents the nutritional condition. Model significance was evaluated using standard linear model summaries and model assumptions regarding residual normality and homoscedasticity were visually inspected and deemed satisfactory. Statistical analyses were carried out using R version 4.2.1 (R Core Team 2024) through the integrated development environment of RStudio (Posit team, 2025), with α = 0.05.

## 3. Results

### 3.1. Biotic and abiotic environmental variables

Table 1 summarizes the mean values ± standard deviation for each environmental variable analyzed, including temperature, salinity, micro- and meso-zooplankton biomasses, N and C composition, isotopic values and C:N ratio. The comparison of the stations where larvae of the three tuna species where captured revealed statistically significant differences for dissolved oxygen concentration, MLD and zooplankton δ¹³C; none of the other environmental variables examined had significant differences. Dissolved-oxygen concentrations were slightly higher (p < 0.05) at stations where ALB was caught than at those where SBT was collected, and were comparable to the values recorded for SKJ. By contrast, the mean MLD associated with SKJ stations was significantly shallower (p < 0.05) than at ALB stations, while showing no difference from SBT stations. δ ¹³C values in both micro- and meso-zooplankton were significantly lower δ¹³C at SKJ stations than at SBT and ALB stations (see Table 1).

**Table 1.** Mean values and ± SE of the environmental variables in the upper 25 m in the study area; parenthesis indicate the number of plankton tows included from which the larvae were collected. Trophic summary of the microzooplankton (55-200 µm) and mesozooplankton (200-2000 µm) size fractions are also shown. Variables were compared with non-parametric Kruskal-Wallis test among SBT, ALB and SKJ species, and Mann-Whitney U test between OPT and DEF growth groups within each species. Different letters indicate significant differences among species (Dunn posthoc test). NS: non-significant, *: p < 0.05, **: p < 0.01

|  | SBT (31) | ALB (15) | SKJ (22) | P |
| --- | --- | --- | --- | --- |
| Environmental variables |  |  |  |  |
| Temperature ($^{\circ}$C) | 28.90 ± 0.07 | 28.70 ± 0.10 | 29.00 ± 0.13 | ns |
| Salinity (ppm) | 34.60 ± 0.03 | 34.50 ± 0.03 | 34.60 ± 0.03 | ns |
| Oxygen (mg l<sup>-1</sup>) | 186.00 ± 3.220<sup>a</sup> | 190.00 ± 0.163<sup>b</sup> | 190.00 ± 0.242<sup>a,b</sup> | * |
| Fluorescence (volts) | 0.107 ± 0.01 | 0.096 ± 0.01 | 0.099 ± 0.00 | ns |
| Mixed Layer Depth (m) | 19.00 ± 2.86<sup>a,b</sup> | 28.20 ± 4.49<sup>b</sup> | 12.70 ± 2.49<sup>a</sup> | * |
| Microzooplakton |  |  |  |  |
| Biomass (mg m<sup>-3</sup>) | 1.12 ± 0.13 | 1.23 ± 0.18 | 0.99 ± 0.15 | ns |
| % N | 6.24 ± 0.18 | 5.97 ± 0.27 | 6.56 ± 0.22 | ns |
| $\delta^{15}$N (‰) | 5.54 ± 0.07 | 5.38 ± 0.10 | 5.66 ± 0.08 | ns |
| %C | 25.87 ± 0.59 | 25.21 ± 0.85 | 25.29 ± 0.70 | ns |
| $\delta^{13}$C (‰) | -18.80 ± 0.10<sup>a,b</sup> | -18.51 ± 0.15<sup>a</sup> | -19.08 ± 0.12<sup>b</sup> | ** |
| C:N | 4.15 ± 0.06 | 4.27 ± 0.09 | 4.03 ± 0.08 | ns |
| Mesozooplaktion |  |  |  |  |
| Biomass (mg m<sup>-3</sup>) | 5.63 ± 0.69 | 6.78 ± 0.98 | 4.98 ± 0.83 | ns |
| % N | 6.96 ± 0.26 | 7.14 ± 0.37 | 7.43 ± 0.30 | ns |
| $\delta^{15}$N (‰) | 6.80 ± 0.08 | 6.54 ± 0.11 | 6.79 ± 0.09 | ns |
| %C | 27.99 ± 0.92 | 28.86 ± 1.30 | 30.78 ± 1.07 | ns |
| $\delta^{13}$C (‰) | -19.02 ± 0.11<sup>a,b</sup> | -18.80 ± 0.15<sup>a</sup> | -19.33 ± 0.13<sup>b</sup> | * |
| C:N | 4.06 ± 0.07 | 4.06 ± 0.11 | 4.19 ± 0.09 | ns |

### 3.2. Larval size and growth

Larval size ranges of SBT, ALB and SKJ are presented in Figure 2A with SBT dominating the spawing grounds while ALB and SKJ were present in more constricted size ranges and in lower abundances. Using the relationships between DW and SL to compare exponential relative growth patterns among the three species, SKJ show the highest weight gain per unit length, while SBT and ALB have faster relative growth in SL (Fig. 2B).

**Figure 2.**
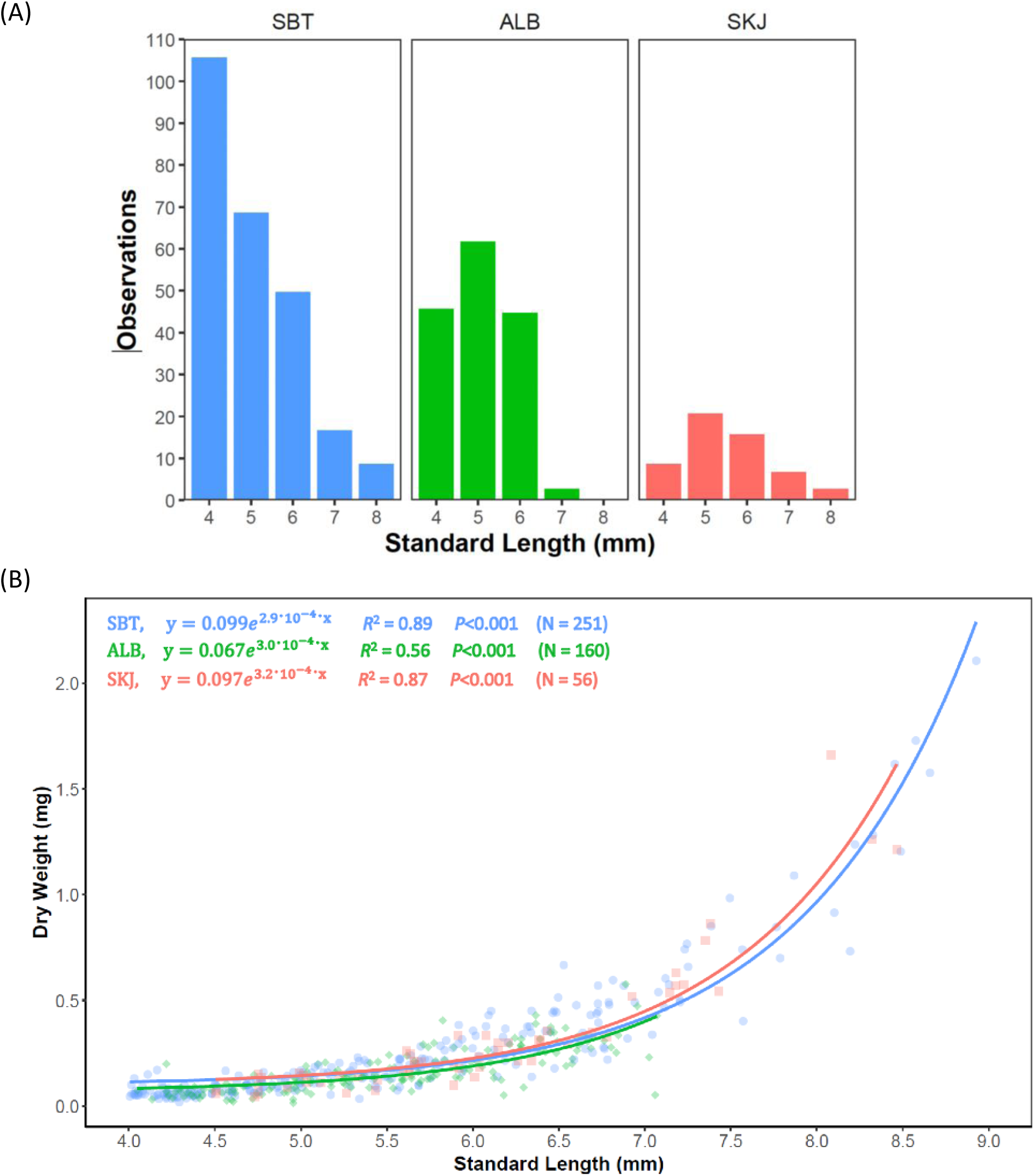
(A) Size (standard length, SL mm) frequency distributions for three tuna species grouped at 1-mm intervals and (B) standard length at dry weight (DW mg). The corresponding least squares regressions in the exponential form (y = a b^x^) are shown for *Thunnus maccoyii* (SBT, blue), *T. alalunga* (ALB, green), and *Katsuwonus pelamis* (SKJ, red) from the eastern Indian Ocean

Larval growth (SL at age) revealed linear but distinct growth trajectories among the three tuna species (Fig. 3A). SBT and ALB exhibited the fastest growth, with slopes of 0.457 and 0.460 mm d^-1^, respectively; while SKJ grew the slowest (0.392 mm d^-1^). All regressions were statistically significant (p < 0.001), with high coefficients of determination (R² > 0.74), indicating strong linear relationships between size and age for each species, and validating that size is a strong predictor of age in these species. The growth rate comparison among species (Fig. 3B), supported these findings and suggest growth variability within each species, with ALB showing the widest variability and exhibiting significantly higher growth rates compared to SBT and SKJ (p < 0.01), whereas no significant difference were observed between SBT and SKJ (Fig. 3B).

**Figure 3.**
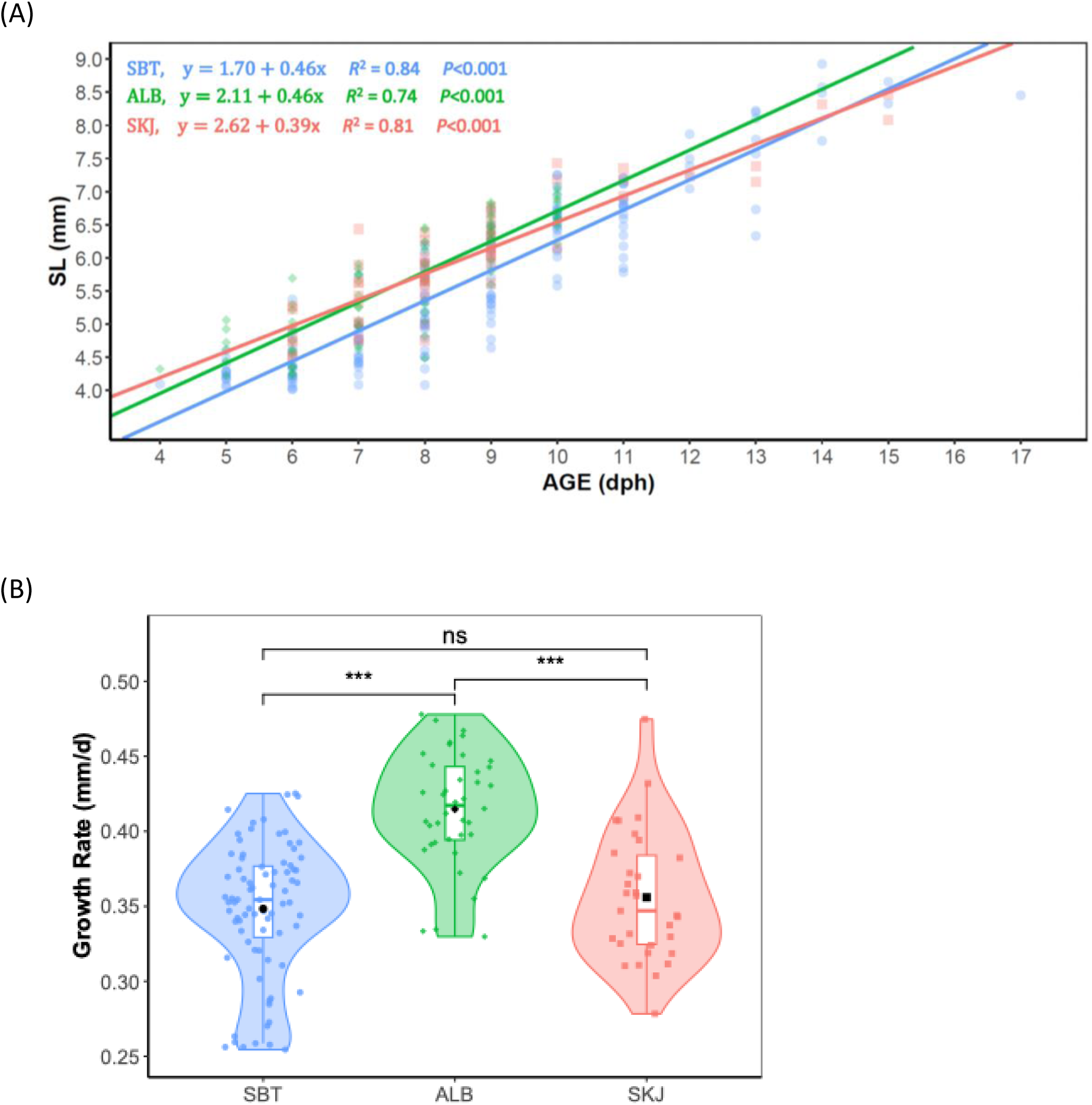
(A) Age (days post-hatch, dph) at standard length (SL, mm) for tuna larvae between 4–9 mm, and (B) box plot within a violin plot of the corresponding growth rates (mm d⁻¹) for the same 4–9 mm size range for *Thunnus maccoyii* (SBT), *T. alalunga* (ALB), and *Katsuwonus pelamis* (SKJ) from the eastern Indian Ocean. Medians are indicated by the horizontal line and statistical differences are indicated at *p* < 0.001 (***).

### 3.3. Larval isotopic analyses

For all larval sizes, the larval ontogenetic relationships for δ^15^N (‰) showed statistically significant (p<0.001) changes in δ^15^N (‰) values over age for SKJ and SBT. The relationships followed a U shape curve, with a sharp decrease until ∼11 and 13 dph for SKJ and SBT, respectively. The curve then increased in δ^15^N values after ∼12 and 16 dph for SKJ and SBT, respectively (Fig. 4A). Higher variability in δ^15^N was observed for younger ages < 8 dph (Fig. 4A). Unlike SKJ and SBT, no significant relationship was observed for ALB (Fig. 4A).

**Figure 4.**
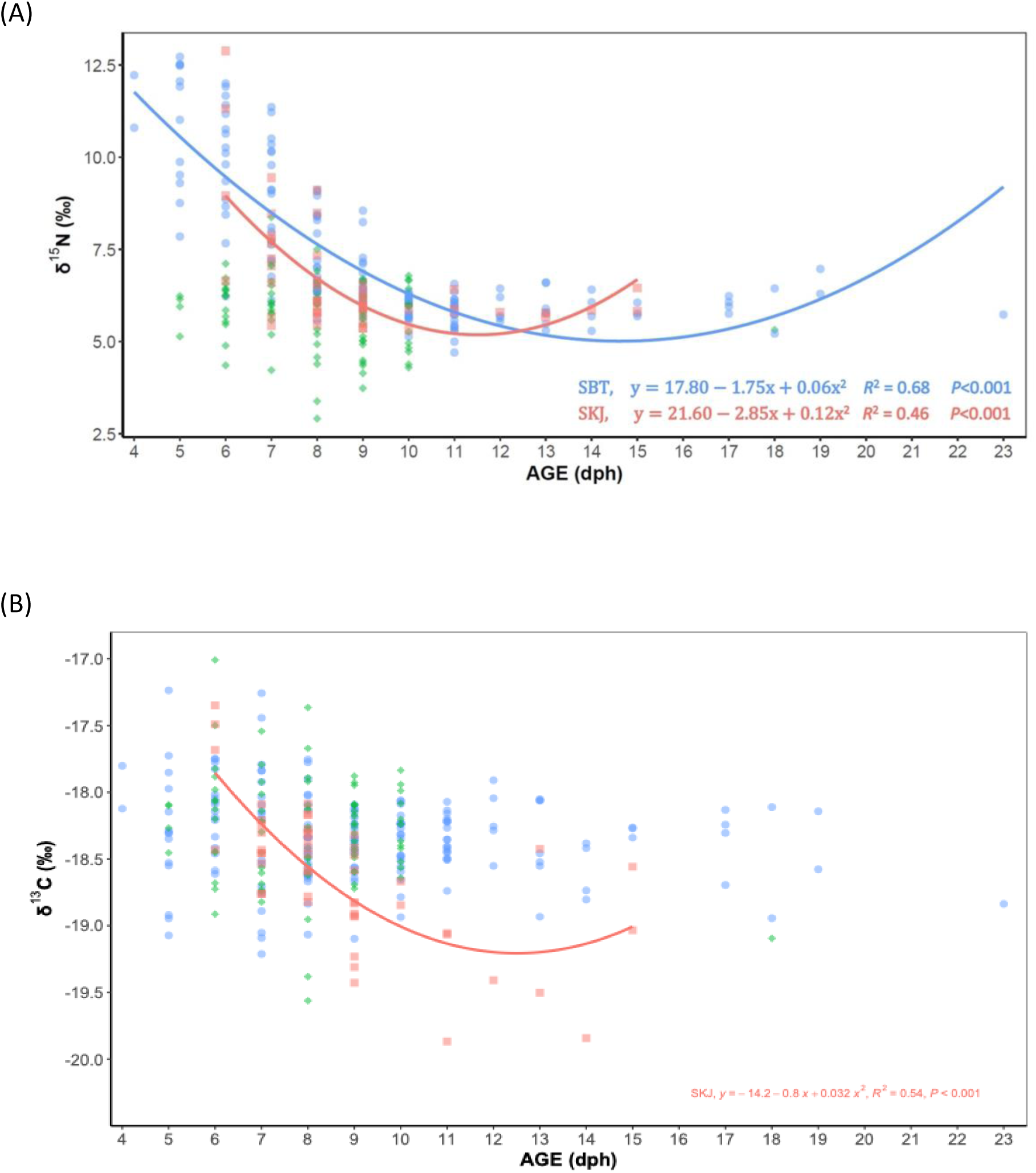
Ontogenetic profiles of (A) δ^15^N and (B) δ^13^C at (AGE, dph) for *Thunnus maccoyii* (SBT), *T. alalunga* (ALB) and *Katsuwonus pelamis* (SKJ) from the eastern Indian Ocean spawning ground. The corresponding least squares regressions in the exponential form (y = a b^x^) are shown for SBT and SKJ.

With respect to δ¹³C (‰) vs age, SKJ showed a significant decreasing relationship with greater variability at younger age until ∼12 dph (p<0.001; Fig. 4B). In contrast, SBT showed no relationship between δ¹³C and age, with δ¹³C appearing relatively stable across all age groups and consistent dispersion of δ¹³C values among ages. No significant relation δ¹³C vs age was observed for ALB (Fig. 4B).

Considering only postflexion larvae, the variation in δ^15^N (‰) over age was weak but statistically significant (p<0.001). A non-linear relationship was observed for SBT, with higher variability observed at younger ages (Fig. 5A). For the δ¹³C (‰) vs age relationship, the regression curves suggested different patterns in δ^13^C values among the species (Fig. 5B).

**Figure 5.**
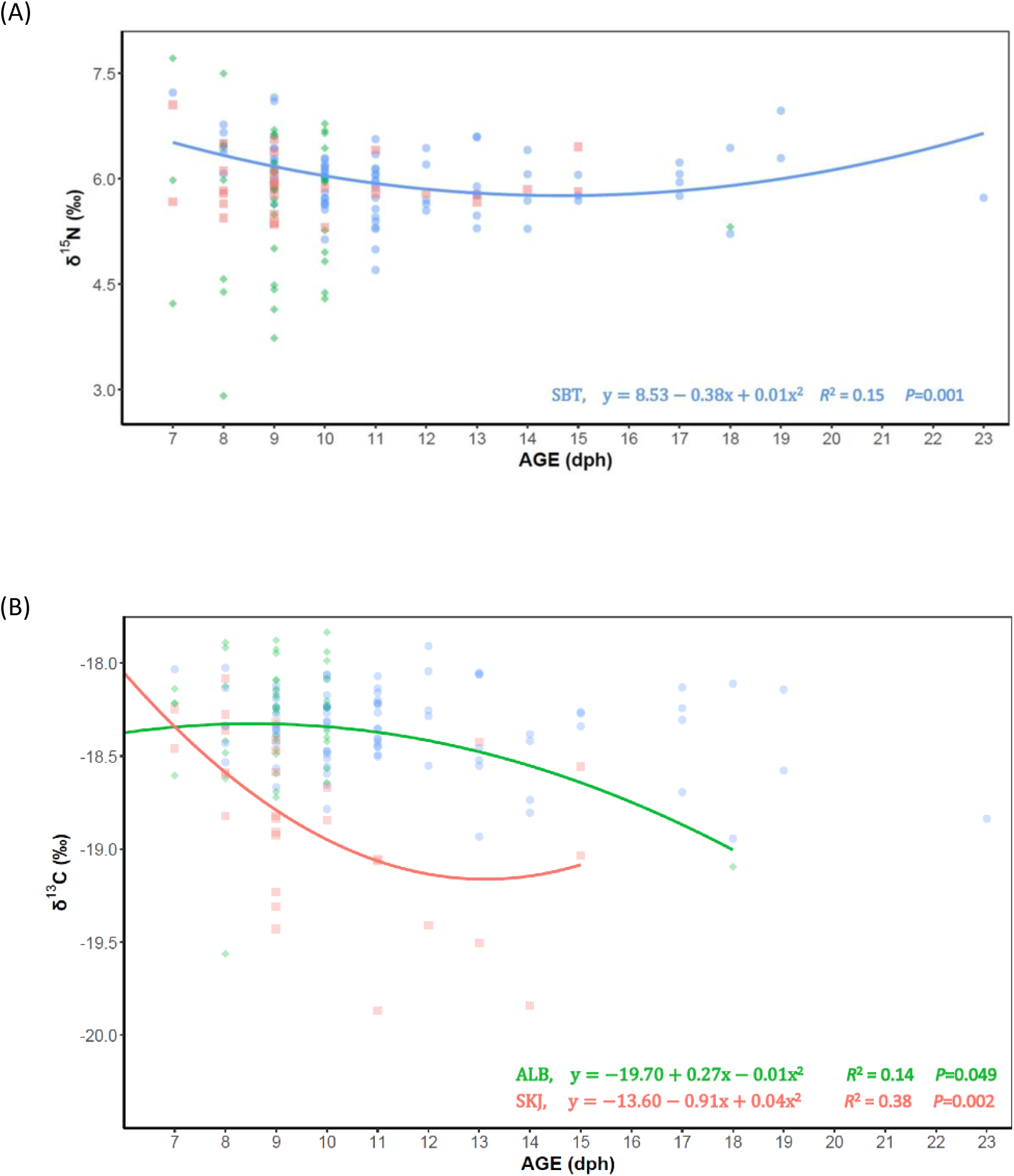
Ontogenetic profiles of (A) δ^15^N and (B) δ^13^C at AGE (dph) for postflexion larval stage for *Thunnus maccoyii* (SBT), *T. alalunga* (ALB) and *Katsuwonus pelamis* (SKJ) from the eastern Indian Ocean spawning ground. The corresponding least squares regressions in the exponential form (y = a b^x^) are shown for SBT, ALB and SKJ.

### 3.4. Larval trophic niches

The isotopic biplots for δ¹⁵N (‰) and δ¹³C (‰) for postflexion SBT, ALB and SKJ indicate species-specific differences in isotopic composition (Fig. 6A). SKJ displays a distinctly depleted δ¹³C signature compared to ALB and SBT, which show overlapping δ¹³C values but are partially separated along their δ¹⁵N signatures. Both the standard ellipse area (SEAc) and bootstrap-based kernel density estimation with rKIN (Figs. 6B, C) consistently show that ALB and SKJ have larger isotopic niches, while SBT exhibits the smallest and most constrained niche. Isotopic niche overlap analysis supports these findings: the highest overlap occurs between SBT and ALB (76% according to SIBER; 93% with KUD), while SKJ shows minimal overlap with both ALB (5% SIBER; 21% KUD) and SBT (12% SIBER; 19% KUD). ALB overlaps moderately with SBT (29% SIBER; 27% KUD), but very little with SKJ (5% SIBER; 14% KUD).

**Figure 6.**
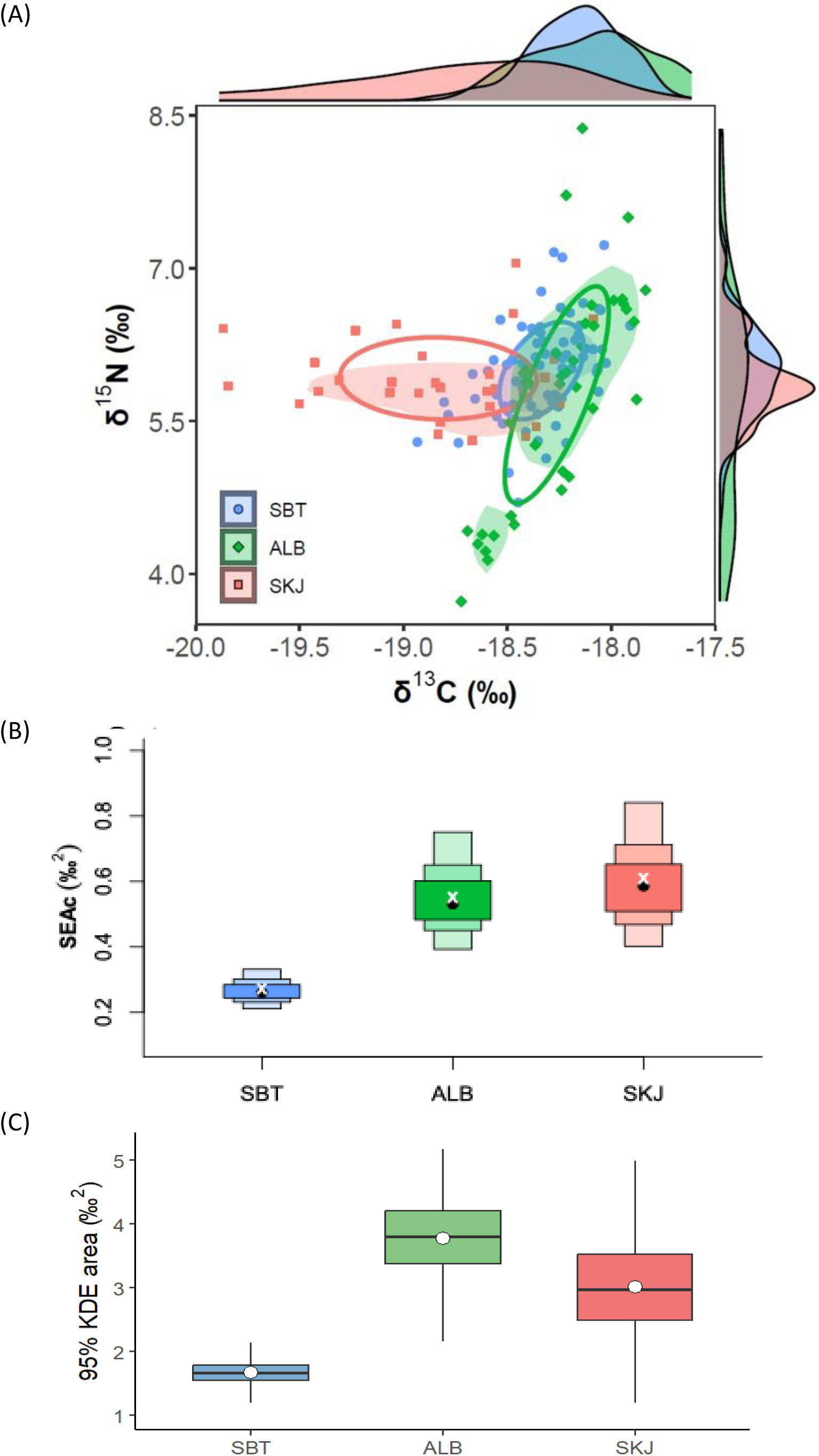
(A) Scatter biplots of δ^13^C and δ^15^N values for postflexion larval stage of *Thunnus maccoyii* (SBT), *T. alalunga* (ALB) and *Katsuwonus pelamis* (SKJ) from the eastern Indian Ocean. Ellipses represent the isotopic niche width for each species, and shaded areas represent the kernel utilization density (KUD, 40% contour). (B) Boxplot of the Bayesian standard ellipse area (SEAc) for the trophic niche of each species. Density plot shows confidence intervals of SEAc corresponding to the mean and 95, 75 and 50% confidence intervals. C) Estimated isotopic niche breadth using rKIN, based on 1000 non-parametric bootstrap iterations per species. For each bootstrap replicate, the 95% kernel density area (KDE) was computed from δ¹³C and δ¹⁵N values. The boxplots represent the distributions of these bootstrap-derived niche areas, with the white point indicating the mean and the box showing the interquartile range.

The intraspecific comparison between residual growth groups (OPT vs DEF), is illustrated for the three studied species in Figure 7, showing their isotopic niche structure for postflexion larvae. In the isotopic biplots, individuals classified as DEF show a broader and more dispersed distribution of δ¹³C and δ¹⁵N values across all three species (Fig. 7A). This visual pattern is confirmed for SBT and ALB by the SEAc-based standard ellipse areas, where DEF groups consistently exhibit wider isotopic niches than OPT individuals, but not for SKJ (Fig. 7B). The bootstrap-based KDE estimates using rKIN reinforce this trend, showing that DEF larvae tend to occupy larger isotopic niche areas, suggesting higher trophic or spatial variability during early development (Fig. 7C). The contrast is most pronounced in ALB, followed by SBT. For SKJ, the OPT and DEF difference is present but more subtle.

**Figure 7.**
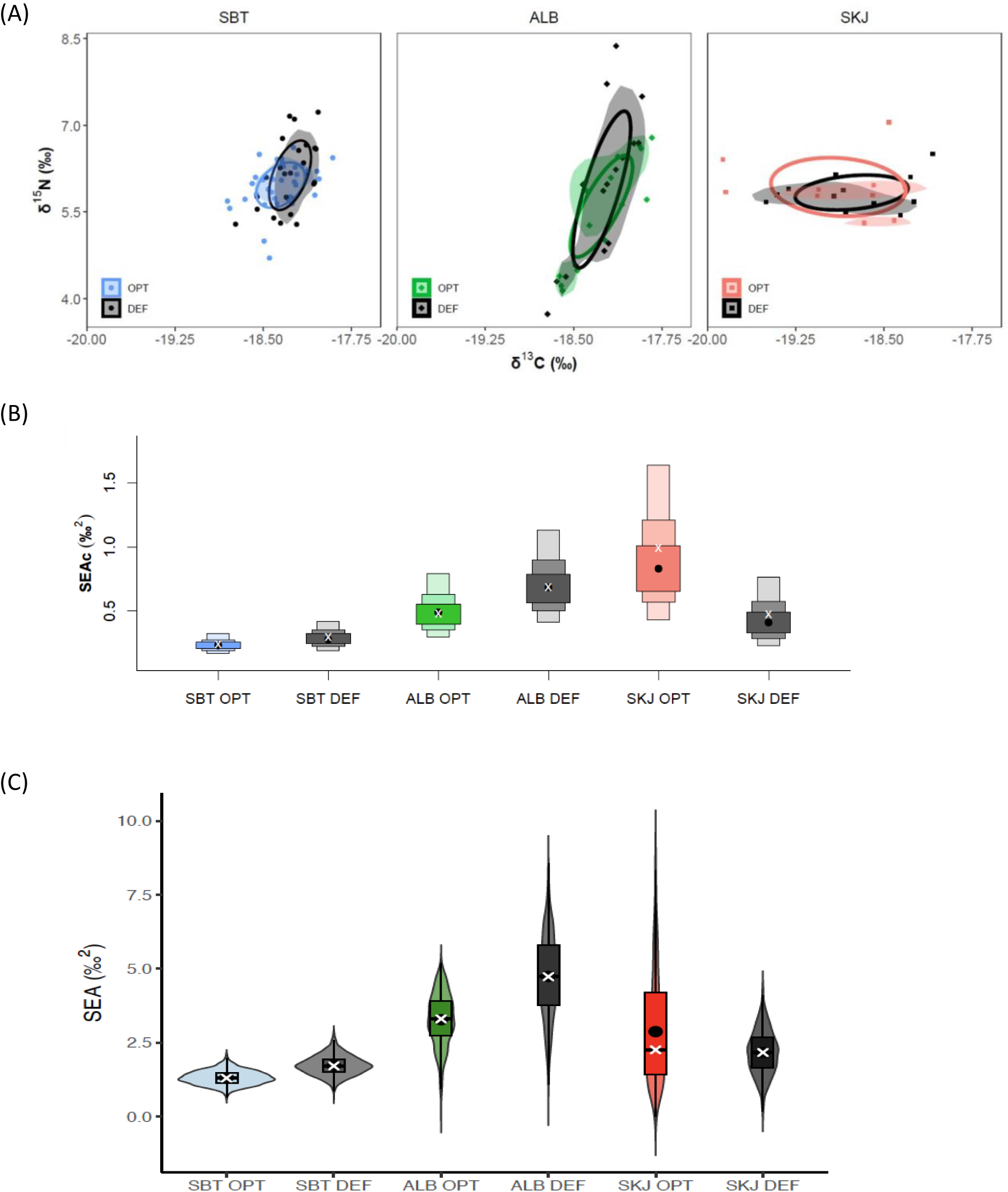
(A) Scatter biplots of δ^13^C and δ^15^N values in OPTIMAL (OPT) and DEFICIENT (DEF) growers for postfelxion larvae of *Thunnus maccoyii* (SBT), *T. alalunga* (ALB) and *Katsuwonus pelamis* (SKJ) from the eastern Indian Ocean. Unfiled ellipses represent the isotopic niche width (SIBER, 40 % of the data regardless of sample size) for each species, and shaded areas represent the kernel utilization density (KUD, 40% contour). B) Boxplot of the corrected standard ellipse area (SEAc) for the trophic niche of each species showing the corresponding confidence intervals for the mean standard ellipse area and 95, 75 and 50%. C) Boxplots within a violin plot for estimated isotopic niche areas using rKIN and 1000 bootstrap replicates. Each iteration involved random resampling with replacement and estimation of the 95% kernel density area (KDE) based on δ¹³C and δ¹⁵N values. The median and mean are indicated by the symbols x and ●, respectively.

### 3.5. Larval trophic positions

The comparison across species indicates that ALB occupies the highest TP, a pattern confirmed by both the frequentist (Fig. 8A) and the Bayesian estimations (Fig. 8B). ALB has the widest spread, (credibility interval, CI: 1.9 – 3.59), with a broader range of TP values than either SKJ (CI: 1.65 – 3.19) or SBT (CI: 1.7 – 3.2), whereas the TP estimates for SBT and SKJ are more tightly clustered. The polynomial model of individual trophic position as a function of age and species revealed statistically significant differences in both intercepts and the linear and quadratic slopes across species (F₈,₁₄₂₃₉₉₁ = 42,330, p < 0.001 for all interaction terms). The species-specific quadratic equations describing the relationship between TP and age, derived from the combined intercepts and slope estimates, are: TP = 4.60 – 0.35 * Age + 0.0132 * Age² for SBT, TP = 7.37 – 0.91 * Age + 0.0437 * Age² for ALB, and TP = 6.86 – 0.86 * Age + 0.0381 * Age² for SKJ. These results indicate that all species exhibit a significant non-linear decline in trophic position with age, with the most pronounced decrease observed in ALB, followed by SKJ, while SBT shows a milder and more stabilized pattern at older ages. The global model fit was highly significant (p < 2.2 × 10⁻¹⁶), and the differences in the shape of the age-individual TP relationship among species reflect distinct ontogenetic dietary shifts. This pattern is consistent with that observed when using standard length and dry weight as predictors (results not shown).

**Figure 8.**
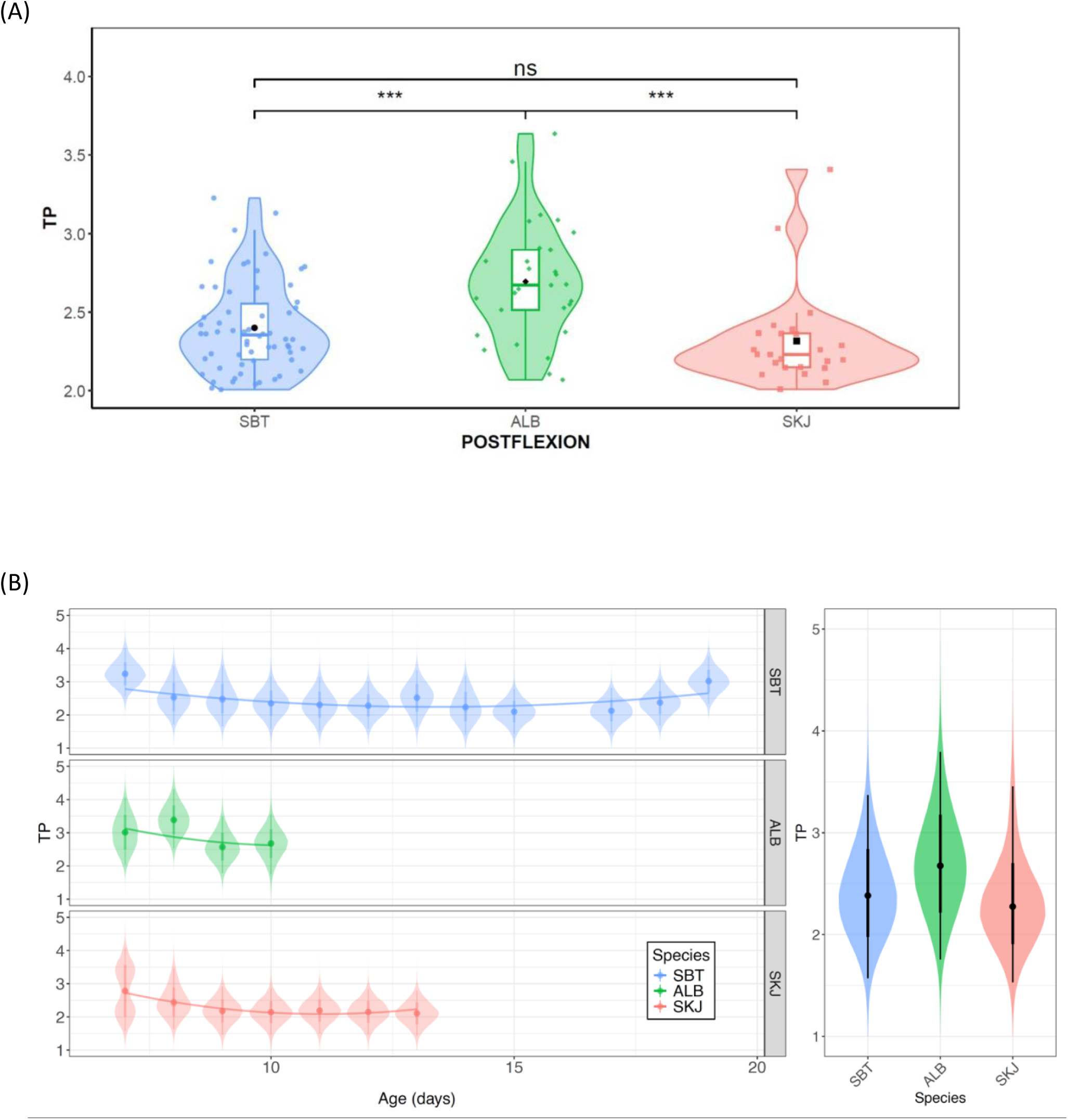
Trophic positions (TP) estimated from (A) Post (2002) model using microzooplankton as primary consumers baseline (TP = 2) and (B) individual Bayesian trophic position estimates in postflexion larval stage of three larval scombrid species: *Thunnus maccoyii* (SBT), *T. alalunga* (ALB) and *Katsuwonus pelamis* (SKJ) from the eastern Indian Ocean common spawning ground. Polynomial models are described in the text. Statistical significance among species is indicated at *p* < 0.001 (***)

Within species, both the frequentist (Fig. 9A) and Bayesian (Fig. 9B) intraspecific analyses reveal that the DEF growth groups for SBT and ALB attain higher TP than the OPT groups. For SKJ, however, these differences were not statistically significant. For the Bayesian analysis, the polynomial models revealed significant differences between OPT and DEF conditions in both the intercept and the age-individual TP relationship across species (p < 0.001 for all terms, complete results not shown). These results indicate that both the magnitude and trajectory of the age-individual TP differ between nutritional conditions and species (Fig. 9B).

**Figure 9.**
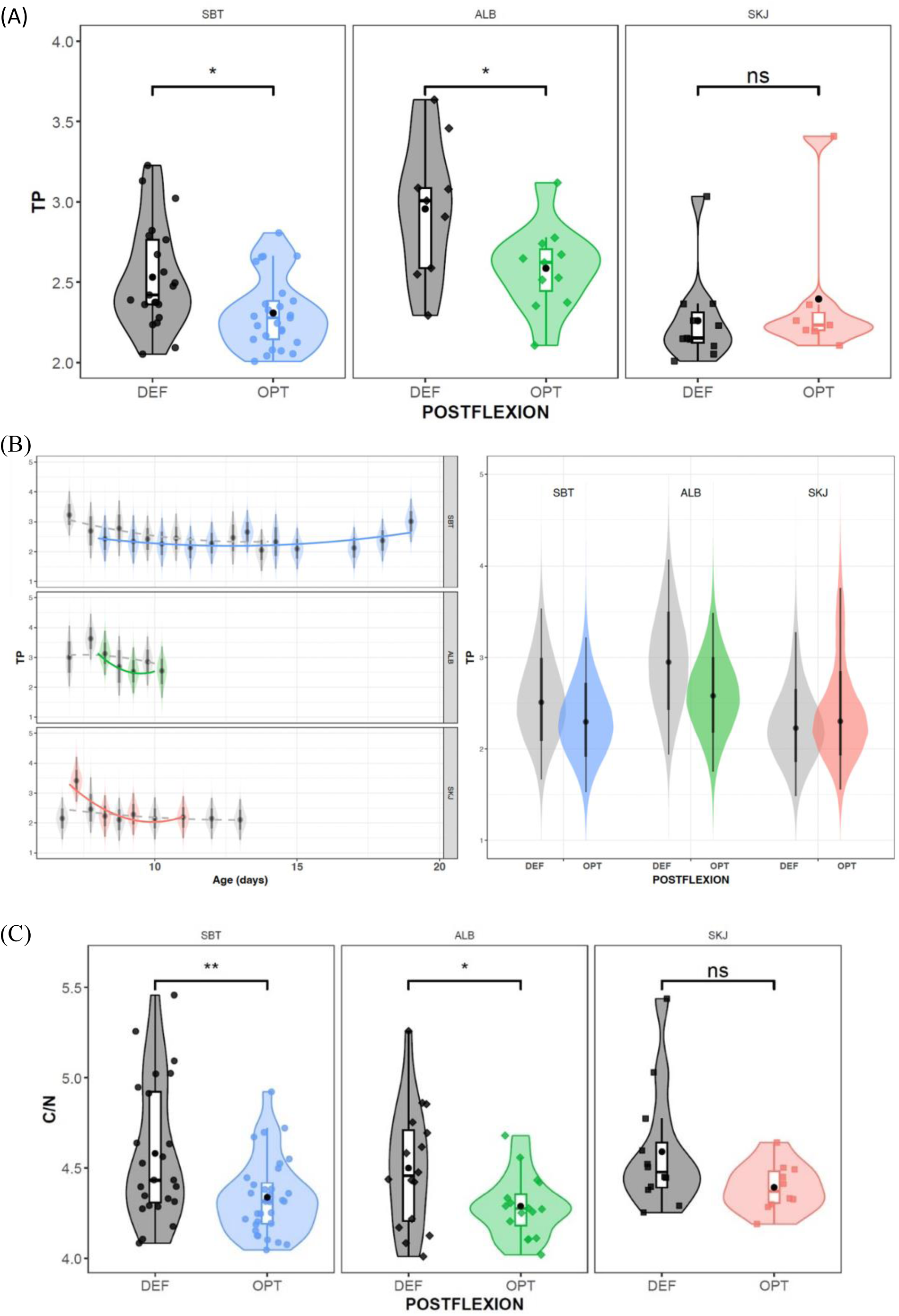
Optimal (OPT) and deficient (DEF) growing groups trophic positions (TP) estimated from (A) Post 2002 model and (B) individual Bayesian model showing the relationship between individual TP and age (in days) for each species. (C) C:N ratio for postflexion larval stages of *Thunnus maccoyii* (SBT), *T. alalunga* (ALB) and *Katsuwonus pelamis* (SKJ) from the eastern Indian Ocean common spawning ground. Statistical significance between the OPT and DEF groups is indicated at the *p* < 0.05 (*) and *p* < 0.01 (**).

### 3.5. Larval nutritional condition (C:N)

No significant differences in C:N ratios has been observed among species, whereas C:N ratios differed significantly between growth groups (DEF and OPT) for ALB and SBT, while a similar, but not significant, trend was observed for SKJ (Fig. 9C). Specifically, DEF larvae exhibited consistently higher C:N values than OPT larvae in ALB (p < 0.05) and SBT (p < 0.01).

## 4. Discusion

Overlap in prey consumption among larval SBT, ALB, and SKJ occurs in our study area (Uotani et al., 1981; Young and Davis 1990) suggesting that while there were similarities in feeding behavior, the specific prey items consumed differ significantly. Our results confirm the same trophic pattern, suggesting shared dietary sources but with clear isotopic niche differentiation and further emphasizing niche segregation between species, with significant larval growth implications. Trophic niche overlap among scombrids suggests competition for resources, although in oligotrophic conditions, partial resource partitioning may mitigate interspecific competition. In our study, faster-growing larvae had significantly narrower isotopic niches, consistent with greater trophic specialization. Furthermore, optimally growing larvae (OPT) had lower TP, suggesting that reliance on efficient, low-level trophic pathways supports enhanced growth performance in the eastern Indian Ocean.

### 4.1. Larval growth

Our results indicate species-specific differences in early larval growth performance. ALB larvae grew faster (i.e., higher growth rates) compared to the other two species, as reflected in both length-at-age and the distributions of daily growth rates. This suggests that ALB may possess intrinsic physiological or ecological traits that enable more efficient somatic growth during the early postflexion stage. In contrast, the lack of significant differences between SBT and SKJ growth rates, despite differences in slope and intercept, implies comparable overall growth performance at similar sizes. García et al. (2006) reported similar growth rates in Mediterranean waters, with fastest ALB larval growth (0.46 mm d^-^¹) during an anomalously warm summer (∼26 °C) in 2003, while growth slowed (0.40 mm d-¹) in cooler summers (∼24 °C) in the same spawning ground (Catalán et al., 2007). Similar larval growth rates were observed for SKJ in the Gulf of México (0.39 mm d^-^¹) in river-influenced waters that seasonally ranged from 22 to 26 °C and in similar upper-mesoscale eddy temperatures observed for co-occurring AFT larvae (Zygas et al., 2015). Additionally, comparable daily growth (∼0.38 mm d^-^¹), were reported at higher temperatures (28–30 °C) in the western-equatorial Pacific, supporting the idea that growth of SKJ plateaus once metabolic requirements are met (Tanabe et al., 2003). However, Uchiyama and Struhsaker (1981) observed slower growth rates (≤ 0.33 mm d⁻¹) in similarly warm water in the central Pacific, a pattern attributed to low prey densities, highlighting the interactive effect of temperature and food supply. Higher temperatures would also require sufficient food to support faster metabolic demands until eventually exceeding the larval termal range (Gleiber et al., 2020).

### 4.2. Ontogenetic isotopic shifts

Marked ontogenetic shifts differ substantially among the three scombrid species examined. U-shaped ontogenetic profiles of δ¹⁵N were observed for SBT and SKJ, with the values sharply dropping in the youngest larvae but increasing again with age, whereas. Similar patterns were previously described for BFT in the Mediterranean Sea (MED, García et al., 2017) and the Gulf of Mexico (GoM, Laiz-Carrión et al., 2019; Quintanilla et al., 2024), where high initial values were attributed to maternally derived isotopic signatures that were gradually diluted as larva growth was fueled by exogenous feeding (Uriarte et al., 2016) on increasingly higher trophic level prey. ALB showed no significant relationship with δ¹⁵N indicating a slower trophic ascent.. These findings agree with earlier regional studies and demonstrate, for the first time, that the maternal isotopic transmission paradigm is not only evident for SBT but also for SKJ. We observe an earlier onset of trophic enrichment consistent with a transition to larger zooplankton prey and its its well-known precocious piscivory (Llopiz et al., 2010; Llopiz and Hobday 2015). SKJ also displayed a significant ontogenetic decline in δ¹³C, consistent with previous studies that suggest a wider vertical habitat distribution with a more diverse diet (Laiz-Carrión et al., 2019). In contrast, SBT maintained an almost stable δ¹³C, and ALB displayed no detectable trend. Our study area is an oligotrophic offshore region with smaller spatial gradients in δ¹³C compared to more coastally influenced regions (i.e., Gulf of Mexico). Overall, SKJ showed stronger isotopic shifts, highlighting a broadening niche and precocious piscivory at much smaller size (∼5 mm in different regions; Llopiz et al., 2010; Young and Davis, 1990; Llopiz and Hobday, 2015) compared with SBT (∼5.4 mm in the IO; Borrego et al., this issue) and ALB (∼ 5.5 mm; Catalán et al., 2007).

### 4.3. Larval trophic niche analyses

Our inter-specific comparisons (Fig. 6A), revealed a clear trophic segregation among the three scombrids. SKJ occupies the most isotopically δ¹³C depleted space, and had minimal overlap with SBT and ALB, that both share a substantial portion of isotopic niche indicating similar carbon sources (Table 2). The narrow SBT niche mirrors the low taxonomic breadth observed through stomach-content analyses for this same cohort (Swalethorp et al., this issue). When combining our SIA with stomach contents analysis, we find support for the hypothesis that SBT exhibit a more selective feeding strategy, potentially conferring advantages for growth and survival. Such traits, combined with distinct reproductive strategies, may help to understand better the predominance of SBT within the ichthyoplankton community during this time of the year (Malca et al., this issue).

**Table 2.** Isotopic niche overlaps of % estimated by standard ellipses corrected areas (SEAc, an ellipse that contains 40% of the data regardless of sample size) and kernel utilization density (KUD, 40% contour) for postflexion larval stages of *Thunnus maccoyii* (SBT), *T. alalunga* (ALB) and *Katsuwonus pelamis* (SKJ) collected in the eastern Indian Ocean spawning ground.

|  | SEAc |  |  | KUD |  |  |
| --- | --- | --- | --- | --- | --- | --- |
|  | *T. macoyii* | *T. alalunga* | *K. pelamis* | *T. macoyii* | *T. alalunga* | *K. pelamis* |
| *T. macoyii* | - | 76 | 26 | - | 94 | 43 |
| *T. alalunga* | 29 | - | 5 | 27 | - | 14 |
| *K. pelamis* | 12 | 5 | - | 19 | 21 | - |

The agreement between isotope- and gut-based metrics strengthens the inference that active prey selection, rather than mere prey encounter, regulates larval growth trajectories in scombrids. Consistent with SKJ broad diet and trophic segregation observed in this study, Laiz-Carrión et al. (2019) reported that SKJ had the largest SEAc among four tuna larvae species examined in the GoM, with a wider δ¹³C range and small overlaps with other tunas. They attributed this breadth to precocious piscivory and mixed carbon sources, conclusions that fit the broad δ¹³C dispersion we observe in the IO. Therefore, intrinsic traits rather than basin-specific responses for SKJ larvae consistently display exceptional trophic plasticity, presumably enabling them to exploit heterogeneous prey fields due to a deeper vertical distributional range (surface - 50 m; (Davis et al., 1990a,b; Llopiz and Hobday, 2015). In contrast, ABT larvae and PBT have been found in shallower depths (upper 20m) (Kodama et al., 2020, Alvarez et al., 2021). The trophic segregation observed at early life stages for these tree scombrid species allows coexistence among them (Varela et al., 2024). Resource partitioning is an adaptative mechanism thant has been previously observed in scombrid larvae (García et al., 2017; Laiz-Carrión et al., 2019) allowing multiple species coexist in the share the same spawning area.

The broader isotopic niche of SKJ coupled with its wide distributional range is consistent consistent with the resource breadth hypothesis (Rader et al., 2017). The hypothesis states that organisms utilizing a wider range of resources can sustain viable populations under a greater variety of environmental conditions, allowing them to occupy larger geographic ranges (Boulangeat et al., 2012). Similarly, Slatyer et al. (2013) found a strong positive relationship between geographic range and both environmental tolerance and habitat use breadth. SKJ has a global spawning distribution and spawn year-round in equatorial and tropical IO waters and seasonally in subtropical waters (Artetxe-Arrate et al., 2021). Furthermore, the broader vertical distribution of SKJ larvae give them access to a more diverse variety of resources compared with SBT and ALB. Similarely, ALB also exhibits a broad isotopic niche and like SKJ it inhabits all tropical, subtropical and temperate pelagic ecosystems, spawning seasonally in subtropical waters of the IO. In contrast, SBT has a constricted spawning season and much more limited geographical spawning range for spawning habitat (Farley and Davis, 1998; Nikoli et al., 2017).

The intra-specific analyses (OPT vs DEF growth groups) reveal a consistent pattern among species: optimally growing larvae (OPT) have smaller isotopic niches, whereas deficient (DEF) larvae occupy broader, more variable niches. These results highlight potential physiological or ecological differences in resource use associated with larval condition and suggests that fast growing individuals are more selective meeting their energetic demands by concentrating on a restricted group of high quality prey (Malca et al., 2022), whereas slower growers rely on a more generalist diet reflecting less efficient foraging or compensatory feeding on lower value prey. These findings align with previous studies emphasizing the role of trophic dynamics in larval growth, survival and recruitment (Pepin et al., 2015; Shropshire et al., 2022; Quintanilla et al., 2025). For both inter- and intra-specific comparisons, the results drawn from both the SIBER and rKIN approaches underscore the robustness of the observed patterns and reveals that this effects of feeding behavior on larval growth variability is consistant across species.

### 4.4. Larval trophic position estimations

Baysian TP estimation demonstrated ALB having a significantly higher and broader TP range than SBT and SKJ which were both very similay (Fig. 7A), with no significant differences between SBT and SKJ. Consistent results appear in applying Bayesian TP estimations (Fig. 7B). These findings agree with Laiz-Carrión et al. (2019), who reported similar TP for ABT than SKJ in the GoM using Compound Specific Stable Isotopes Analyses of Amino Acids (CSIA-AA). In the MED, García et al. (2017) found that ALB larvae occupied a slightly narrower isotopic niche than ABT, but they did not detect significant TP divergence once baseline variability was accounted for. Instead, ALB showed a wide δ¹⁵N range similar to that observed in our study. Further analyses of TP based on CSIA-AA may elucidate these inter-specific differences accounting their baselines adaptability. In this study we report the first TP estimates for postflexion SBT larvae which were (2.51 ± 0.32) compared with similarly sized ABT larvae in the GoM (2.76 ± 0.32 from Laiz Carrion et al., 2019; 4.38 ± 0.29 and 3.65 ± 0.24 from Quintanilla et al., 2024; and 3.74 ± 0.59 from Malca et al., 2023) and in the MED (3.46 ± 0.59; Malca et al., 2023). Differences in δ¹⁵N baseline, including interannual variability, could explain these differences between water basins. However, the distinct biologies, trophic preferences, metabolic requirements and habitat adaptations of different bluefin tuna poulations coupled with factors such as food availability and interactions with other species could significantly influence estimates of their TPs. In a complementary analysis from the BLOOFINZ study, the diet of SBT larvae was shown to primarily consist of appendicularians that through utilization of the microbial food web would result in a short and efficient food chain for the larvae (Stukel et al., 2022; Swalethorp et al, this issue).

In the present analysis, faster-growing larvae occupied markedly narrower isotopic niches, a pattern consistent with greater trophic specialization. Furthermore, the DEF (slow-growing) groups of SBT and ALB exhibited lower mean TP, whereas SKJ showed no clear difference between growth groups. DEF larvae also displayed higher TP variability among-individuals, suggesting that larvae with reduced growth rates exploit a broader and less consistent array of food resources. These patterns should be interpreted cautiously, however, because the number of postflexion larvae available after residual-growth analyses was limited, especially for SKJ. Our findings parallel those of Quintanilla et al. (2024), who reported that ABT larvae in the Gulf of Mexico grew fastest when their TP was lowest accross two consecutive years. A similar relationship has been documented for larval Shortbelly Rockfish, where individuals with lower TP were heavier and grew more rapidly (Kwan et al., 2024). Together, these studies and our results also suggests that TP is indicative of the amount of energy reaching the reaching the larval population (Montoya, 2007; Caut et al., 2009) with a low TP equal high energy transfer efficiency from the base of the food chain up to the larvae supporting higher performance as is stipulated in the trophic efficiency in early life hypothesis (Swalethorp et al., 2023). Applying this rationale to our data, SBT and ALB larvae would achieve optimal growth when they rely on efficient, low-level trophic pathways, where quantity and/or nutritional quality of diet may be optimal to sustain higher growth potential and survival within the larval population (Swalethorp et al., 2023).

### 4.5. Larval nutritional condition

The C:N ratio serves as a biochemical proxy for the nutritional condition of fish larvae, particularly reflecting their lipid reserves. Higher C:N values typically indicates greater lipid content, while lower ratios suggest lipid depletion due to energy investment in somatic growth (Logan et al., 2008; Kloppmann et al., 2022). In our analysis, C:N ratios differed between growth groups, with DEF larvae exhibiting consistently higher C:N ratios compared to their OPT counterparts, indicating a reduced utilization of lipid reserves (and potentially reduced muscle mass buildup).. These findings align with previous observations in ABT, where higher C:N values in DEF larvae were interpreted as a reflection of lower energetic investment in growth and reduced mobilization of lipid stores (Quintanilla et al., 2024). In contrast, OPT larvae across species generally exhibited lower C:N values, consistent with elevated growth rates and a corresponding reduction in lipid content due to increased metabolic demand. An additional none-excluding interpretation is that a higher C:N ratio may reflect a lower proportion of structural protein relative to OPT individuals, which is consistent with what would be expected in DEF larvae. These intra-specific patterns reinforce the use of C:N as a reliable indicator of larval physiological condition that can be interpreted alongside growth rates as a consequence of a higher growth potential. They consequently support the broader hypothesis that growth potential is closely tied to energy allocation strategies during early ontogeny

## 5. Conclusions and perspectives

To our knowledge, this study represents the first comprehensive application of stable isotope analysis to study the trophic ecology of SBT larvae. Our findings provide valuable temporally integrated insights into the TP and trophic niche of SBT, corroborating specific dietary patterns identified from stomach content analysis (Swalethorp et al., this issue). These results highlight the utility of integrated dietary approaches in detecting ontogenetic shifts in prey consumption and resource partitioning during early developmental stages. This holds relevance not only for SBT, but also for other ecologically and commercially important species such as ALB and SKJ. The patterns we observed may reflect species-specific life history strategies or environmental adaptations that drive daily growth variability. Such interspecific differences in growth have important implications for larval survival, recruitment success, and overall population dynamics, particularly in ecosystems where growth rate is closely tied to predation risk and competitive fitness.

The concordance between isotope- and gut-based metrics strengthens the inference that active prey selection, rather than mere prey encounter, regulates larval growth trajectories in scombrids. Upcoming research should apply amino acid compound specific isotopic analysis and metabarcoding techniques to further resolve dietary selectivity and refine TP estimates for both larvae and their prey, linked with daily growth variability.

Estimates of tuna larvae abundance are used as indices of spawning stock abundance in stock assessments that are the basis of management advice. For most tunas, however, the relationship between spawning stock abundance and recruitment remains uncertain. This study has contributed to improving our understanding of why this may be so paving the way to developing less uncertain stock assessment advice. Ultimately, a better understanding of larval ecosystem dynamics will allow for more robust management strategies fulfilling the principles of ecosystem-based fishery management.

## Acknowledgements

We deeply thank A. García, J. Lamkin and T. Gerard for opening and aligning our collaboration groups from Spain and USA. We sincerely thank L.E. Beckley, A. Jivanjee and L. Mattison during the tuna larvae sampling and processing onboard, extending our gratitude to the crew of the *R/V Roger Revelle* for their assistance above and beyond during the RR2201 cruise. This study was funded by the INDITUN project PID2021/122862NB/100 UE-FEDER (R.L.-C.) and grant PRX23/00189 (R.L.-C.) supported by the Ministry of Science, Innovation and Universities (MICINN) of the Spanish government, U. S. National Science Foundation awards BLOOFINZ-IO OCE-1851558 (M.R.L.) and OCE-1851558 (D.D. and E.M.), and contributes to the Second International Indian Ocean Expedition (IIOE-2 endorsed project EP046). Zooplankton and larval fish samples were collected under Australian Government permit AU-COM2021-520 and Australian Marine Parks permit PA2021-00062-2 issued by the Director of National Parks, Australia. Views expressed in this publication do not necessarily represent those of the Director of National Parks or the Australian Government. Special thanks to the R development community for providing essential tools for data analysis. R.L.-C. gratefully acknowledges financial support by the Fulbright Program, which is sponsored by the U.S. Department of State, the U.S. – Spain Fulbright Commission.

## Author Statement

R.L.-C, J.M.Q., R.B.-S., D.D. R.S.and M.R.L. conceived the study. R.L.-C, J.M.Q. and E.M., collected and/or analyzed zooplankton samples; R.L.-C, J.M.Q., R.B.-S. and M.A.G., collected and/or analyzed larval fish samples; J.M.Q. and R.B.-S. analyzed otolith microstructures; R.L.-C., M.A.G., and R.B.-S. processed the samples for SIA. R.L.-C, J.M.Q., R.B.-S, C.Q.-R. R.S. and F.J.A. analyzed the data and discuss de results. R.L.-C, R.S. and E.M. wrote the manuscript. All authors provided comments and edits on the manuscript.

## Declaration of generative AI and AI-assisted technologies in the writing process

Generative AI and AI-assisted technologies were solely employed during the writing process to improve the readability and language clarity of the manuscript. After using this tool/service, the author(s) reviewed and edited the content as needed and take(s) full responsibility for the content of the publication.

